# Cholinergic boutons are distributed along the dendrites and somata of VIP neurons in the inferior colliculus

**DOI:** 10.1101/2022.09.18.508423

**Authors:** Julia T. Kwapiszewski, Luis M. Rivera-Perez, Michael T. Roberts

**Author notes:** **Correspondence:** Michael T. Roberts.

## Abstract

Cholinergic signaling shapes sound processing and plasticity in the inferior colliculus (IC), the midbrain hub of the central auditory system, but how cholinergic terminals contact and influence individual neuron types in the IC remains largely unknown. Using pharmacology and electrophysiology, we recently found that acetylcholine strongly excites VIP neurons, a class of glutamatergic principal neurons in the IC, by activating α_3_β_4_* nicotinic acetylcholine receptors (nAChRs). Here, we confirm and extend these results using tissue from mice of both sexes. First, we show that mRNA encoding α_3_ and β_4_ nAChR subunits is expressed in many neurons throughout the IC, including most VIP neurons, suggesting that these subunits, which are rare in the brain, are important mediators of cholinergic signaling in the IC.

Next, by combining fluorescent labeling of VIP neurons and immunofluorescence against the vesicular acetylcholine transporter (VAChT), we show that individual VIP neurons in the central nucleus of the IC (ICc) are contacted by a large number of cholinergic boutons. Cholinergic boutons were distributed adjacent to the somata and along the full length of the dendritic arbors of VIP neurons, positioning cholinergic signaling to affect synaptic computations arising throughout the somatodendritic compartments of VIP neurons. In addition, cholinergic boutons were observed in close apposition to dendritic spines on VIP neurons, raising the possibility that cholinergic signaling also modulates presynaptic release onto VIP neurons. Together, these results strengthen evidence that cholinergic signaling exerts widespread influence on auditory computations performed by VIP neurons and other neurons in the IC.

## Introduction

The inferior colliculus (IC), the midbrain nucleus of the central auditory system, receives extensive cholinergic input from the pedunculopontine tegmental nucleus (PPT) and the laterodorsal tegmental nucleus (LDT) (Motts and Schofield 2009). Activity in cholinergic PPT and LDT neurons is influenced by sensory novelty, arousal, rewards, and the sleep-wake cycle, providing a rich repertoire of behavioral conditions that might drive cholinergic modulation of sound processing in the IC (Reese et al. 1995a; Reese et al. 1995b; Sakai 2012; Boucetta et al. 2014; Van Dort et al. 2015; Gut and Winn 2016; Kroeger et al. 2017). Consistent with this, cholinergic signaling in the IC can decrease stimulus specific adaptation (Ayala and Malmierca 2015), sharpen frequency tuning (Askew et al. 2017), enhance temporal processing (Felix et al. 2019), and promote cortically driven plasticity (Ji et al. 2001; Ji and Suga 2009). It is likely that additional functions for cholinergic signaling in the IC remain to be discovered.

Anatomical evidence also suggests that cholinergic signaling strongly influences computations in the IC. Cholinergic axons from the PPT extend throughout the three major subdivisions of the IC (central nucleus, dorsal cortex, and lateral cortex), where they form a dense network of presumptive release sites, and routinely contact the somata of glutamatergic and GABAergic IC neurons (Noftz et al. 2020). Cholinergic boutons are also routinely found adjacent to the proximal dendrites of many glutamatergic and GABAergic IC neurons, including GABAergic neurons from each of the four subtypes defined in the IC based on soma size and extracellular markers (Beebe and Schofield 2021). Complimenting this, IC neurons express some of the highest levels of muscarinic and nicotinic receptors found in the brain (Cortes et al. 1984; Clarke et al. 1985; Schwartz 1986; Glendenning and Baker 1988; Wada et al. 1989; Morley and Happe 2000; Whiteaker et al. 2002; Gahring et al. 2004; Happe and Morley 2004; Sottile et al. 2017). However, despite multiple lines of evidence pointing to the importance of cholinergic signaling in the IC, how cholinergic terminals contact and influence molecularly defined classes of IC neurons remains largely unknown.

We recently showed that VIP neurons in the IC are strongly excited by acetylcholine (Rivera-Perez et al. 2021). VIP neurons are a molecularly defined class of glutamatergic stellate neurons found throughout the major subdivisions of the IC. Because VIP neurons project to multiple targets, including contralateral IC, auditory thalamus (medial geniculate body), superior colliculus, periaqueductal gray, and ventral nucleus of the trapezoid body (Goyer et al. 2019), cholinergic modulation of VIP neurons has the potential to impact auditory and non-auditory computations in multiple brain regions. Using pharmacology and patch clamp recordings in brain slices from mice, we found that acetylcholine excites VIP neurons primarily by activating α_3_β_4_* nicotinic acetylcholine receptors (nAChRs; * *indicates that the identity of the fifth subunit in the receptor pentamer is unknown*), with a minor contribution from α_7_ nAChRs. We also provided qualitative evidence that cholinergic boutons could be found closely apposed to the somata and dendrites of VIP neurons.

In the present study, we confirm and extend these previous findings. First, using single molecule fluorescence in situ hybridization, we show that most VIP neurons, and many other IC neurons, express mRNA for the α_3_ and β_4_ nAChR subunits. Next, using immunofluorescence against the vesicular acetylcholine transporter (VAChT), we found that cholinergic boutons were routinely located in close apposition to the somata and dendrites of VIP neurons in the central nucleus of the IC (ICc), including along the full length of the dendrites of VIP neurons that were reconstructed based on biocytin fills. Quantification revealed that the dendritic arbors of each fully reconstructed VIP neuron were contacted by >100 cholinergic boutons and that these boutons were evenly distributed along the proximal to distal axis of the dendrites. Finally, we found that cholinergic boutons were also often located adjacent to the dendritic spines of VIP neurons, where they might either directly synapse onto VIP neurons or influence the release of glutamate onto VIP neurons. Together, these results suggest that cholinergic signaling can influence synaptic computations throughout the somatic and dendritic compartments of VIP neurons.

## Materials and Methods

### Animals

All experiments were approved by the University of Michigan Institutional Animal Care and Use Committee and were in accordance with NIH guidelines for the care and use of laboratory animals. Mice were kept on a 12-h day/night cycle with *ad libitum* access to food and water. VIP-IRES-Cre mice (*Vip^tm1(cre)Zjh^*/J, Jackson Laboratory, stock # 010908) (Taniguchi et al. 2011) were crossed with Ai14 reporter mice (*B6.Cg-Gt(ROSA)26Sor^tm14(CAG-tdTomato)Hze^*/J, Jackson Laboratory, stock #007914)(Madisen et al. 2010) to yield F1 offspring that expressed the fluorescent protein tdTomato in VIP neurons. F1 mice of both sexes, aged P30 – P85, were used for experiments.

### Fluorescence in situ hybridization

Fluorescence in situ hybridization was performed on fresh frozen brain tissue from three VIP-IRES-Cre × Ai14 mice (one P47 female and two P58 males) using the RNAScope Multiplex Fluorescent V2 Assay (Advanced Cell Diagnostics, Newark, CA). To prepare fresh frozen brain tissue, mice were deeply anesthetized with isoflurane and transcardially perfused with phosphate buffered saline (PBS). Brains were then rapidly removed by dissection, frozen on dry ice, embedded in cryo-embedding medium, and frozen at −80°C. Cryo-embedded brains were subsequently sectioned using a cryostat at −20°C to obtain 15 μm-thick coronal sections of the IC. IC sections were placed on microscope slides and stored at −80°C. The RNAScope Assay was run according to the manufacturer’s protocol using probes targeting mRNA encoding the α_3_ nAChR subunit (*Chrna3*, Channel 1 probe), the β_4_ nAChR subunit (*Chrnb4*, Channel 2 probe), and *tdTomato* (Channel 3 probe). In brief, the RNAScope Assay consisted of: Tissue sections were fixed in 10% neutral buffered formalin and dehydrated using a series of 50%, 70%, and 100% ethanol. Sections were then treated with hydrogen peroxide for 10 minutes at room temperature and then with Protease IV for 30 minutes at room temperature. Probes were hybridized and the hybridize amplification steps were run as follows AMP 1 for: 30 minutes at 40°C, AMP 2 for 30 minutes at 40°C, AMP 3 for 15 minutes at 40°C. To develop the Channel 1 signal, HRP-C1 was applied for 15 minutes at 40°C, and Opal 520 for 30 minutes at 40°C. The same steps were followed for Channels 2 and 3, with Opal 570 used for Channel 2 and Opal 690 for Channel 3. Opal dyes were obtained from Akoya Biosciences (Marlborough, MA). Tissue was counterstained with DAPI for 30 seconds at room temperature, a coverslip was placed over the sections, and the sections were left to dry overnight. Slides were stored at 4°C and imaged within 2 weeks after the protocol was finished. Tile-scan images of complete IC sections were collected using a 20× objective and 0.5 μm Z-step size on a Leica TCS SP8 laser scanning confocal microscope. Images of the slices were analyzed using Neurolucida 360 (MBF Bioscience, Williston, VT).

### Detection of cholinergic terminals adjacent to VIP neuron somata and dendritic segments in the ICc

VIP-IRES-Cre × Ai14 mice of both sexes aged P30-P85 were deeply anesthetized with isoflurane and transcardially perfused with PBS for 30 seconds and then with a 10% neutral buffered formalin solution (Sigma-Aldrich, catalog # HT501128, St. Louis, MO) for 15 minutes. Brains were removed, post-fixed in the same formalin solution for 2 hours, washed in PBS, and then cryoprotected overnight at 4°C in antifreeze solution containing 2.5 parts PBS, 1.5 parts ethylene glycol, and 1 part glycerol (volume/volume). Brains were then sliced into 40 μm coronal sections on a Leica VT1200S vibrating microtome. Sections containing the IC were stored at −20°C in antifreeze solution for no more than 30 days. Slices were removed from the freezer and washed in PBS, then treated for 2 hours in a PBS solution containing 0.3% TritonX-100 and 10% normal donkey serum (NDS, Jackson ImmunoResearch Laboratories, West Grove, PA). Sections were then treated overnight at 4°C with a PBS solution containing 1% NDS, 0.3% TritonX-100, and rabbit anti-VAChT (3:500, Synaptic Systems, catalog # 139103, RRID: AB_887864, Goettingen, Germany). The VAChT antibody was previously validated in a VAChT-knockout study (Kolisnyk et al. 2013) and has been used to identify cholinergic terminals in the cochlear nucleus, the intercollicular midbrain, and hippocampus (Goyer et al. 2016; Gillet et al. 2018; Zhang et al. 2019; Noftz et al. 2021). The next day, slices were washed in PBS and then treated for 2 hours at room temperature with PBS with 0.3% TritonX-100, 1% NDS, and Alexa Fluor 647-tagged donkey anti-rabbit IgG (1:500, Thermo Fisher, catalog # A-31573, RRID: AB_2536183, Waltham, MA). Slices were then washed in PBS, mounted on gelatin-subbed slides (SouthernBiotech, catalog # SLD01-BX, Birmingham, AL), and coverslipped using Fluoromount-G mounting medium (SouthernBiotech, catalog # 0100–01).

Slices were imaged on a Leica TCS SP8 laser scanning confocal microscope. For each analyzed section, a tile-scan image of one side of the IC was collected using a 20× objective. The borders of the ICc were delineated following the approach developed by Choy Buentello et al. (2015), for which we used an atlas of anti-GAD67 and anti-GlyT2-stained IC sections prepared for a previous study from our lab (Anair et al. 2022). Regions of interest from the ICc that contained one or more VIP neuron somata were then chosen for analysis by examining only the tdTomato fluorescence channel (i.e., the VIP neuron channel); immunofluorescence from the VAChT channel was not visualized during the neuron selection process to avoid biasing results. Z-stack images of the selected regions were collected using a 63×, 1.40 NA oil-immersion objective and 0.1 μm Z-steps. Once collected, VIP neuron somata and dendrites in the region of interest, including dendritic segments not connected to a soma in the image stack, were reconstructed using Neurolucida 360. VAChT+ puncta adjacent to reconstructed VIP neuron somata or dendrites were then identified using a two-step process in Neurolucida 360. First, we used Neurolucida’s automatic puncta detection algorithm with a 5 μm detector diameter and 70% detector sensitivity to detect puncta that were <2 μm from a reconstructed soma or dendrite and had a volume >500 voxels (0.4 μm^3^). Second, we used a manual curation process to correct for detected puncta that were obviously mislabeled (e.g., cases where a punctum was clearly much larger than an axonal swelling or where multiple adjacent puncta had been counted as one). Data were then imported to Neurolucida Explorer (MBF Bioscience), where puncta analysis tools were used to calculate total puncta, puncta around soma, puncta around dendrite, length of dendrite, average nearest neighbor, closest nearest neighbor, average puncta volume, smallest puncta volume, and largest puncta volume. The closest nearest neighbor and volume measures were used to check for outlier cases, which when detected, were then put through the manual curation process again.

### Detection of cholinergic terminals adjacent to fully reconstructed VIP neurons in the ICc

Whole-cell patch-clamp recordings were targeted to VIP neurons in the ICc of acutely prepared brain slices from VIP-IRES-Cre × Ai14 mice aged P45 – P80. Mice were deeply anesthetized with isoflurane and then rapidly decapitated. The brain was removed, and a tissue block containing the IC was dissected in 34°C ACSF containing the following (in mM): 125 NaCl, 12.5 glucose, 25 NaHCO_3_, 3 KCl, 1.25 NaH_2_PO_4_, 1.5 CaCl_2_ and 1 MgSO_4_, bubbled to a pH of 7.4 with 5% CO_2_ in 95% O_2_. Coronal sections of the IC (200 μm) were cut in 34°C ACSF with a vibrating microtome (VT1200S, Leica Biosystems) and incubated at 34°C for 30 min in ACSF bubbled with 5% CO_2_ in 95% O_2_. Slices were then incubated at room temperature for at least 30 min before being transferred to the recording chamber. Slices were placed in a recording chamber under a fixed stage upright microscope (BX51WI, Olympus Life Sciences) and were constantly perfused with 34°C ACSF at ~2 ml/min. Electrodes were pulled from borosilicate glass capillaries (outer diameter 1.5 mm, inner diameter 0.86 mm, Sutter Instrument, Novato, CA) to a resistance of 3.5 – 5.0 MΩ using a P-1000 microelectrode puller (Sutter Instrument). The electrode internal solution contained (in mM): 115 K-gluconate, 7.73 KCl, 0.5 EGTA, 10 HEPES, 10 Na_2_ phosphocreatine, 4 MgATP, 0.3 NaGTP, supplemented with 0.1% biocytin (w/v), pH adjusted to 7.4 with KOH and osmolality to 290 mmol/kg with sucrose. After 10 minutes of recording, the electrode was carefully moved away from the neuron and out of the bath. Slices were immediately placed in formalin for 24 hours and moved to fresh PBS the next day for immunofluorescence treatment.

Fixed slices were rinsed in PBS and then treated with 10% NDS and 0.3% TritonX-100 for 2 hours. Slices were then incubated for 48 hours at 4°C in PBS with 0.3% TritonX-100, 10% NDS, rabbit anti-VAChT (1:500, Synaptic Systems, catalog # 139103, RRID: AB_887864), and streptavidin-Alexa Fluor 488 (1:1000, Thermo Fisher, catalog # S11223). Sections were next rinsed in PBS, treated with 10% NDS and 0.3% TritonX-100 in PBS for 30 minutes, and then incubated in Alexa Fluor 647-tagged donkey antirabbit IgG (1:500, Thermo Fisher, catalog # A-31573, RRID: AB_2536183) for 24 hours at 4°C. Sections were then mounted on slides and coverslipped using Fluoromount-G. Z-stack images of streptavidin-Alexa Fluor 488-stained neurons were collected using a 1.40 NA 63× oil-immersion objective and 0.1 μm Z-steps on a Leica TCS SP8 laser scanning confocal microscope.

Images were imported into Neurolucida 360 software (MBF Bioscience) where the somata and dendritic arbors of biocytin-filled VIP neurons were reconstructed. After reconstruction, the distribution of VAChT+ puncta adjacent to reconstructed neurons was determined following the same procedure detailed above except that detected puncta had to be <1 μm from the dendrites or soma of the reconstructed cell and have a volume >75 voxels (0.06 μm^3^).

Sholl analysis was conducted using Neuolucida Explorer on the fully reconstructed neurons and their detected adjacent puncta. The Sholl analysis generated concentric spheres centered around the centroid of the cell body with radii increasing in 10 μm increments. The 10 μm shells were used to quantify the number of puncta and the length of dendrite within each shell. To account for the variable density of dendrites at varying lengths away from the cell body, the puncta were then normalized to the number of puncta per 100 μm of dendrite. The first 10 μm shell primarily contained somata, with little or no dendritic length, and was therefore excluded from subsequent analysis.

## Results

Here, we report results from experiments performed on brain tissue from VIP-IRES-Cre × Ai14 mice of both sexes. VIP neurons were identified by the expression of the fluorescent protein tdTomato, as previously described (Goyer et al. 2019).

### Most VIP neurons and many other IC neurons express mRNA encoding α_3_ and β_4_ nAChR subunits

Although α_3_ and β_4_ nAChR subunits have limited expression in the brain, radioligand-binding and in situ-hybridization studies indicate that the IC is one of the few brain regions where these subunits are expressed (Wada et al. 1989; Whiteaker et al. 2002; Marks et al. 2002; Salas et al. 2003; Gahring et al. 2004; Marks et al. 2006). Consistent with this, we recently found that excitation of VIP neurons by acetylcholine is largely blocked by SR16584, an antagonist selective for α_3_β_4_* nAChRs (Rivera-Perez et al. 2021). Since the specificity of receptor antagonists is rarely perfect, especially for antagonists against receptors that can exist in many heteromeric combinations, we tested whether these pharmacological results were supported by the expression of α_3_ and β_4_ subunits in VIP neurons. To do this, we performed single molecule fluorescence in situ hybridization (RNAScope) on coronal sections from three VIP-IRES–Cre × Ai14 mice (one P47 female and two P58 males) using probes against *CHRNA3* (mRNA encoding the α_3_ subunit), *CHRNB4* (mRNA encoding the β_4_ subunit), and *tdTomato* (mRNA used to identify VIP neurons). This experiment also allowed us to test whether other neurons in the IC, in addition to VIP neurons, express α_3_ and β_4_ nAChR subunits.

Our results showed that *CHRNA3* (cyan) and *CHRNB4* (yellow) were widely expressed throughout the three major subdivisions of the IC (**Figure 1A**; ICc, dorsal cortex, and lateral cortex). VIP neurons, identified by *tdTomato* expression (magenta), were also distributed throughout the three major IC subdivisions, consistent with our past results (**Figure 1A**) (Goyer et al. 2019). Strong punctate labeling for *CHRNA3* and *CHRNB4* was detected in many IC neurons, including in VIP neurons (magenta; **Figure 1B,C**). To quantify nAChR-subunit expression in VIP neurons, we examined 8 representative IC sections spread across the rostral-caudal axis of the IC from a total of 3 mice. All *tdTomato*-expressing cells found in these sections, regardless of IC subdivision, were marked in Neurolucida 360 and then examined for expression of *CHRNA3* and *CHRNB4*. Among VIP neurons, 98% expressed *CHRNA3* (cyan; 366 of 373 *tdTomato+* cells), 61% expressed *CHRNB4* (yellow; 227 of 373 *tdTomato+* cells), and 61% expressed both *CHRNA3* and *CHRNB4* mRNA (226 of 373 *tdTomato+* cells). Thus, most VIP neurons have the potential to express α_3_β_4_* nAChRs. Interestingly, while all but one VIP neuron that expressed *CHRNB4* also expressed *CHRNA3*, 37% of VIP neurons expressed *CHRNA3* but not *CHRNB4*. This suggests that a portion of VIP neurons expresses nAChRs containing α_3_ subunits combined with β subunits not of the β_4_ type.

**Figure 1.**
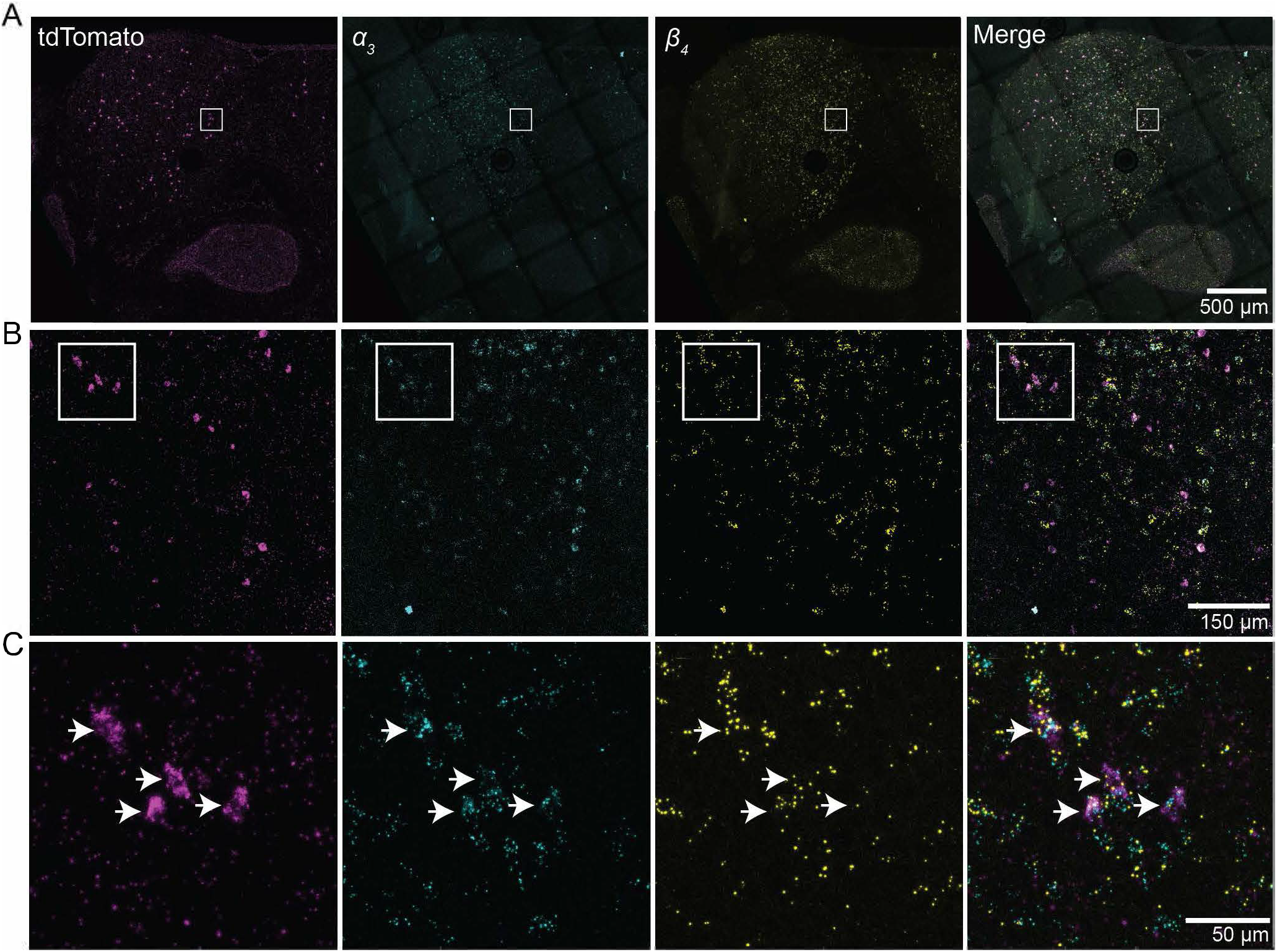
VIP neurons and many other IC neurons express mRNA for α_3_ and β_4_ nAChR subunits. Single molecule in situ hybridization (RNAScope) was performed on coronal sections of the IC with probes against *tdTomato, CHRNA3* (α_3_ nAChR), and *CHRNB4* (β_4_ nAChR) mRNAs. **A)** Low magnification tile-scan images of the left side of a coronal section of the IC showing labeling with probes against *tdTomato* (left, magenta), *CHRNA3* (second from left, cyan), and *CHRNB4* (third from left, yellow), with a merged image shown on the right. **B)** Intermediate magnification images of the region of interest indicated by the boxes in **A**. **C)** High magnification images of the region of interest indicated by the boxes in **B**. Arrows highlight three example cells that labeled with all three probes, indicating that VIP neurons (tdTomato+) often express mRNA encoding α_3_ and β_4_ nAChR subunits. *CHRNA3* and *CHRNB4* mRNA were also routinely co-localized in cells that were negative for tdTomato, indicating that expression of mRNA encoding α_3_ and β_4_ nAChR subunits is also common among other neuron types in the IC.

Expression of *CHRNA3* and/or *CHRNB4* was also routinely observed among non-VIP cells (*tdTomato*-) in the IC (**Figure 1C**; cells identified by DAPI staining, DAPI channel not shown). In non-VIP cells, *CHRNA3* and *CHRNB4* puncta were commonly co-expressed but could also be expressed individually. These results support the hypothesis that α_3_- and β_4_-containing nAChRs are important mediators of cholinergic responses in the IC.

### Distribution of VAChT+ puncta adjacent to VIP neurons in the ICc

Although VIP neurons express nAChR subunits (**Figure 1**) and depolarize in response to exogenous applications of acetylcholine (Rivera-Perez et al. 2021), it is not clear whether cholinergic input to VIP neurons is common or how cholinergic inputs are distributed along the somatic and dendritic compartments of VIP neurons. These are important questions to address because the anatomical arrangement of cholinergic input to VIP neurons will constrain hypotheses about how cholinergic signaling influences VIP neurons. To assess the distribution of putative contacts between cholinergic projections and VIP neurons in the ICc, we used VAChT immunofluorescence to identify cholinergic projections in IC sections prepared from VIP-IRES-Cre × Ai14 mice. VAChT immunofluorescence was observed throughout the IC (**Figure 2A**), similar to a recent report detailing VAChT immunofluorescence in intercollicular areas in rats (Noftz et al. 2021). VIP neurons, which were identified by the expression of tdTomato (**Figure 2B**), were also observed in the major IC subdivisions and were nearly always in regions rich with VAChT immunofluorescence (**Figure 2C**).

**Figure 2.**
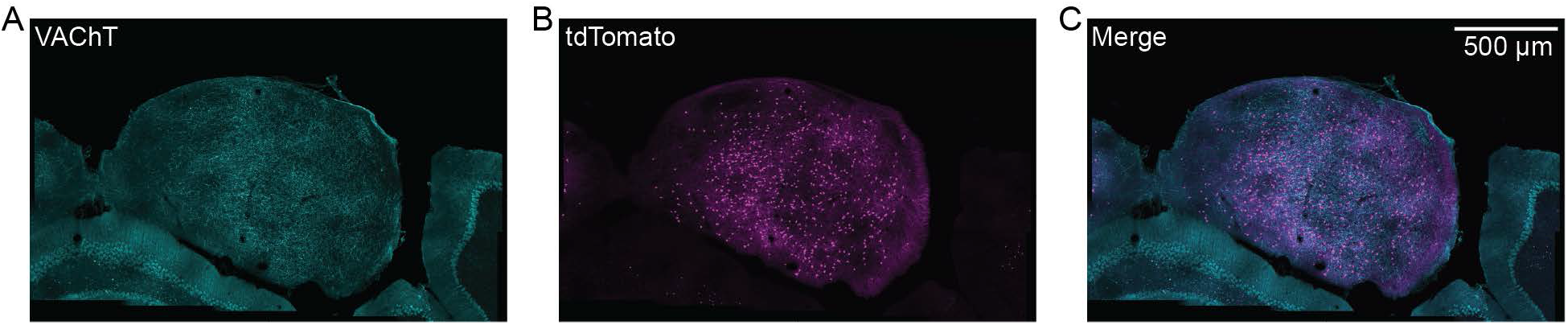
Cholinergic projections and VIP neurons had overlapping distributions in the IC. Low magnification images of a caudal section of the IC show that VAChT+ projections (**A**, cyan) and VIP neurons (**B**, magenta) were broadly distributed in the IC and generally had strongly overlapping distributions (**C**, merge). Scale bar in **C** applies to all panels.

To test for potential contact sites between cholinergic boutons or terminals and VIP neurons in the ICc, we examined VAChT-stained IC sections from VIP-IRES-Cre × Ai14 mice and identified regions of interest within the ICc that contained one or more VIP neuron (tdTomato+) somata. These regions of interest were selected by examining the tdTomato channel alone; the VAChT channel was not visible during selection to avoid biasing results. We then collected high resolution confocal image stacks of the anti-VAChT immunofluorescence and tdTomato fluorescence in the identified regions of interest using a 63x, 1.40 NA oil-immersion objective and 0.1 μm Z-step size. In total, images were collected from 17 regions of interest representing 13 IC sections from 7 mice. In these images, VAChT+ puncta were routinely observed in close contact with VIP neuron somata and dendrites. **Figure 3** shows examples of VAChT+ puncta localized against the surfaces of six different VIP neuron somata, and **Figure 4** shows examples of VAChT+ puncta in close apposition to VIP neuron dendritic segments from six regions of interest.

**Figure 3.**
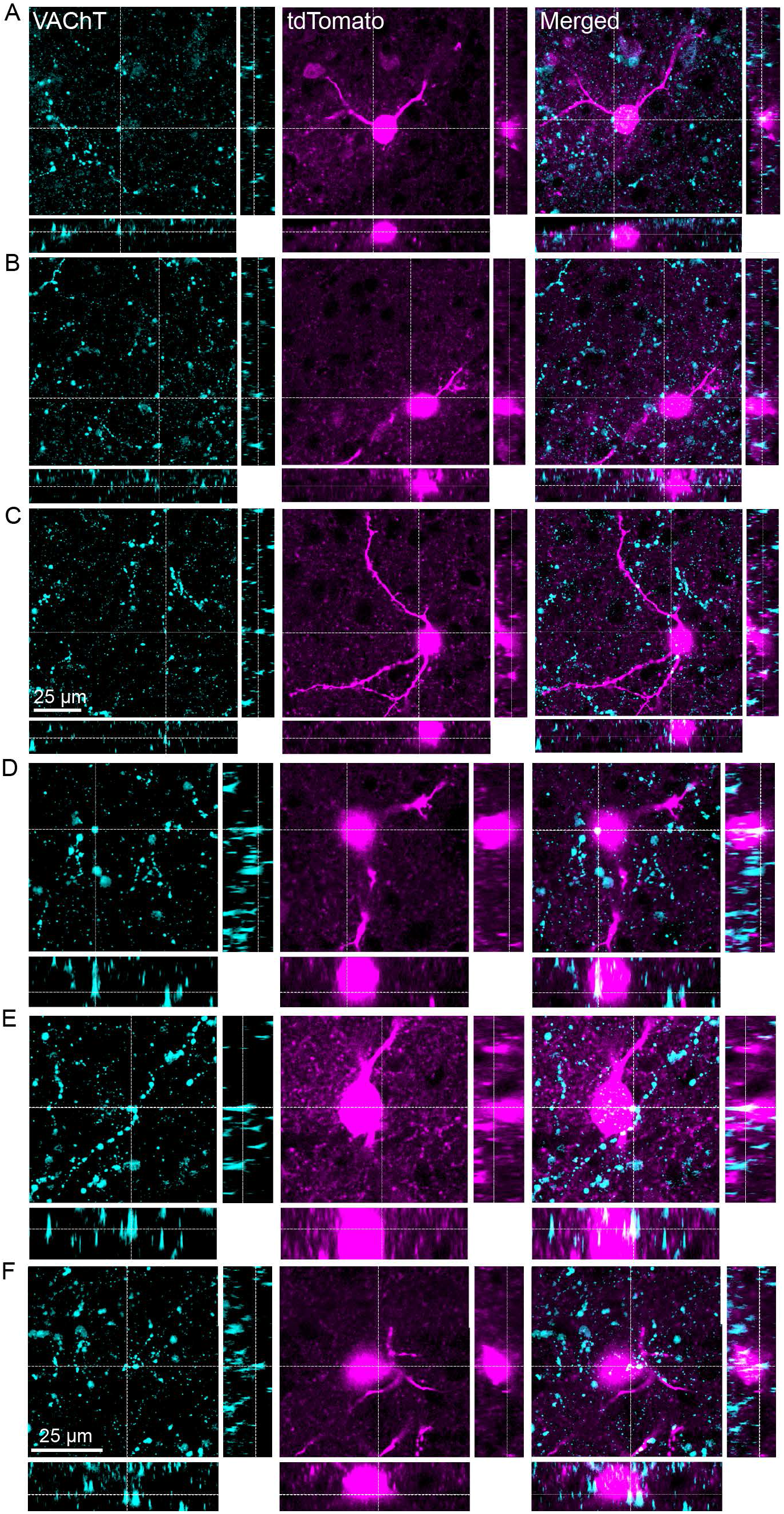
Cholinergic puncta were routinely observed in close apposition to VIP neuron somata. **A-F** show examples of VAChT+ puncta located directly adjacent to the somata of six different VIP neurons. VAChT immunofluorescence is shown in the left column in cyan, tdTomato fluorescence from VIP neurons is shown in the middle column in magenta, and merged images are shown in the right column. Each image stack is shown from three perspectives. The large image represents a maximum intensity projection of the Z-stack as viewed from above. A VAChT+ punctum is highlighted at the intersection of the vertical and horizontal dashed lines. The two smaller images provide side views of virtual optical sections prepared from the Z-stack at the locations indicated by the dashed lines in the large image. The scale bar in **C** applies to **A-C**, and the scale bar in **F** applied to **D-F**. Data are from 4 mice.

**Figure 4.**
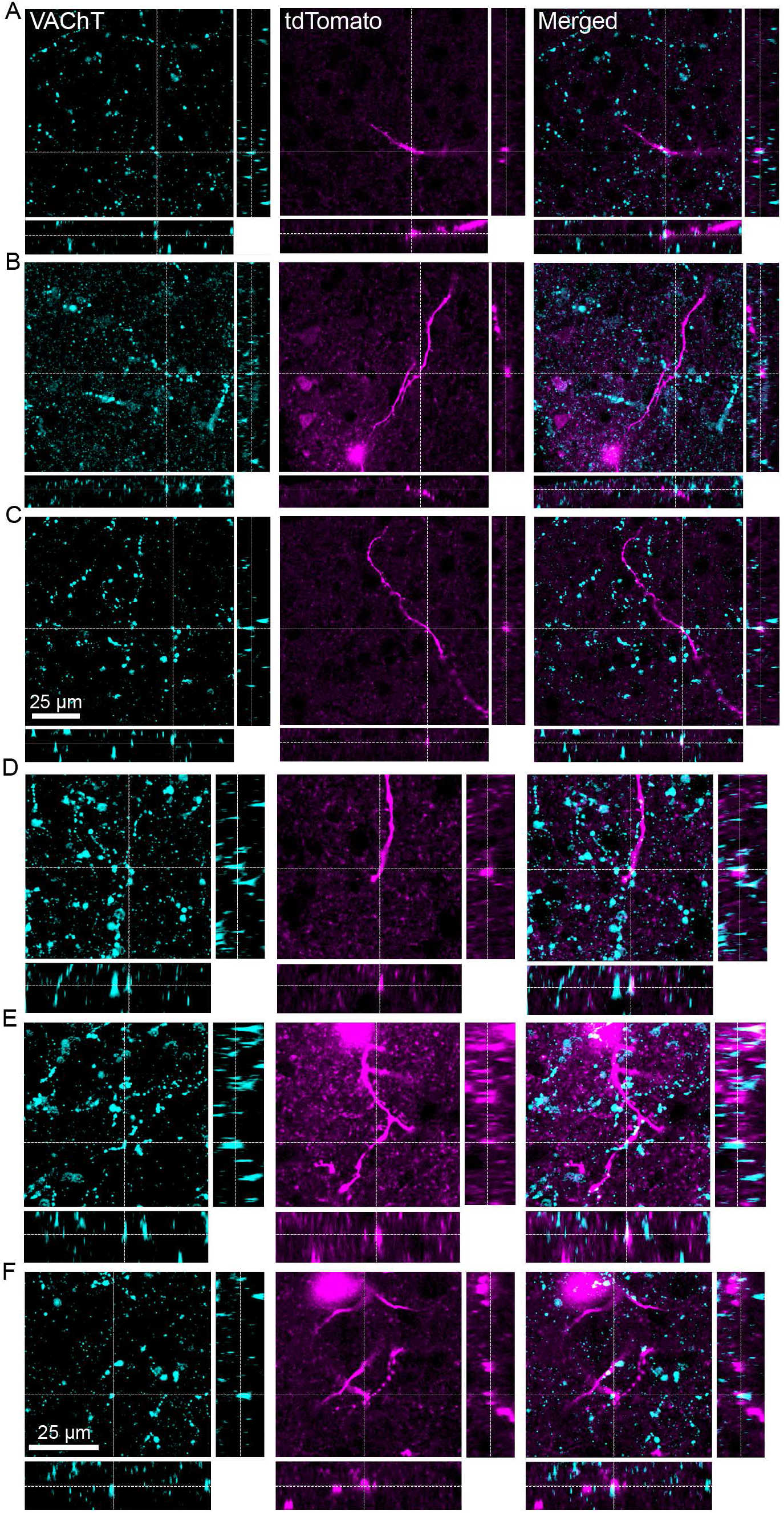
Cholinergic puncta were routinely observed in close apposition to VIP neuron dendrites. **A-F** show examples of VAChT+ puncta located directly adjacent to dendritic segments identified in six different regions of interest. The color scheme and organization of the figure is the same as described in the Figure 3 legend. The scale bar in **C** applies to **A-C**, and the scale bar in **F** applied to **D-F**. Data are from 4 mice.

To quantify the number of potential contact sites between VIP neurons and VAChT+ puncta, we used the puncta detection feature in Neurolucida 360. This involved first reconstructing the VIP neuron somata and dendritic segments in each region of interest and then using the puncta detection algorithm to detect VAChT+ puncta that were <2 μm from a reconstructed soma or dendrite and had a volume >500 voxels (>0.4 μm^3^). We then reviewed the automatically detected puncta and removed rare instances where the algorithm counted multiple puncta as one or marked a punctum as being much larger than was warranted by the underlying VAChT immunofluorescence. The resulting data sets were then imported into Neurolucida Explorer, where results were quantified. This analysis revealed that VIP neuron somata were contacted by an average of 12 ± 5 VAChT+ puncta (mean ± SD; range of 5 – 22), while VIP neuron dendrite segments were contacted by an average of 46 ± 29 VAChT+ puncta (mean ± SD; range of 2 – 101) (**Figure 5A**). Since the analyzed dendritic segments varied in length, we also normalized the dendritic puncta counts per 100 μm of dendrite length. This revealed that VIP neuron dendrites were contacted by an average of 12 ± 8 VAChT+ puncta per 100 μm of dendrite (mean ± SD; range of 2 – 35) (**Figure 5B**). These numbers suggest that putative contacts between cholinergic axons and both the somata and dendrites of VIP neurons in the ICc are quite common.

**Figure 5.**
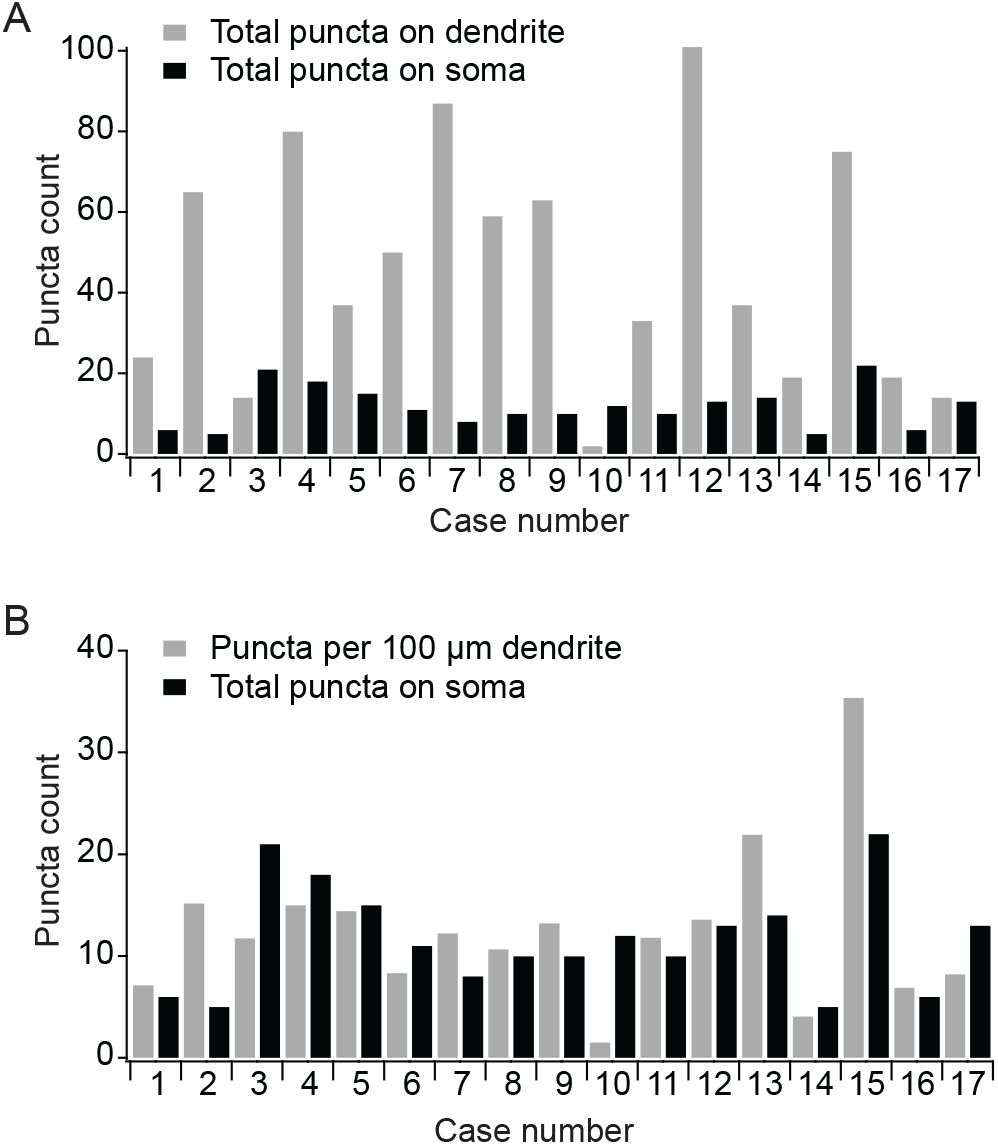
Multiple cholinergic puncta were found in close apposition to each VIP neuron soma and dendritic segment. VIP neuron somata and dendritic segments were reconstructed in Neurolucida 360, and a puncta detection algorithm was used to detect VAChT+ puncta located within 2 μm of each reconstructed element. **A)** Total VAChT+ puncta counts for the somata and dendritic segments reconstructed from 17 regions of interest. Each region of interest contained only one VIP neuron soma. **B)** Same data as **A** but the dendritic puncta counts were normalized to 100 μm of dendritic length to facilitate comparisons across regions of interest, which varied in the total dendritic length they contained.

### Distribution of VAChT+ puncta adjacent to fully reconstructed VIP neurons in the ICc

We next sought to determine the distribution of cholinergic puncta adjacent to the full dendritic arbors of VIP neurons in the ICc, as opposed to the dendritic segments that were analyzed above. To do this, we used whole-cell patch-clamp recordings to fill individual VIP neurons in the ICc of acutely prepared IC slices with biocytin. This allowed for subsequent reconstruction of the full somatodendritic compartments of single VIP neurons. We confirmed that each recorded neuron had a sustained firing pattern and intrinsic physiology consistent with the physiological properties previously described for VIP neurons (Goyer et al. 2019). To preserve full dendritic arbors and enable whole-cell recordings, the brain slices used for this experiment were thicker than those used above (200 μm compared to 40 μm). Presumably due to this change, we found that a slight modification to the anti-VAChT immunofluorescence protocol was necessary to ensure even labeling of VAChT+ puncta throughout the thickness of the slice (see Methods for details). We also found that VAChT+ puncta detection was more accurate in thicker tissue sections when the detection criteria were modified to detect puncta within 1 μm of the reconstructed neuron and to require a minimal puncta volume of 75 voxels (0.06 μm^3^).

The resulting data revealed that VIP neurons in the ICc were closely apposed by putative VAChT+ terminals throughout the entire extent of their dendritic arbors (**Figure 6**). Quantitative analysis showed that the somata of the 9 fully reconstructed VIP neurons were closely apposed by an average of 28 ± 7 VAChT+ puncta (mean ± SD; range of 12 – 39), while the dendritic arbors were lined with an average of 248 ± 97 VAChT+ puncta (mean ± SD; range of 139 – 473; **Figure 7A**). The total length of dendrites reconstructed per neuron in this data set ranged from 1040 – 3230 μm, with a mean ± SD of 1450 ± 700 μm. Normalizing the VAChT+ puncta counts for dendrite length revealed that the VIP neuron dendrites in the fully reconstructed dataset were contacted by an average of 17 ± 4 puncta per 100 μm of dendrite (mean ± SD; range of 12 – 22; **Figure 7B**). This result is similar to the 12 ± 8 VAChT+ puncta per 100 μm of dendrite observed for the isolated dendritic segments above. Interestingly, there was no correlation in the fully reconstructed neurons between the VAChT+ puncta per 100 μm dendrite and the total dendritic length per neuron (Pearson’s *r* = −0.39, p = 0.29). Thus, puncta density did not systematically change as a function of dendritic arbor size. This is also visually apparent in **Figure 7**, where the reconstruction cases are arranged by total dendritic length, from shortest on the left to longest on the right.

**Figure 6.**
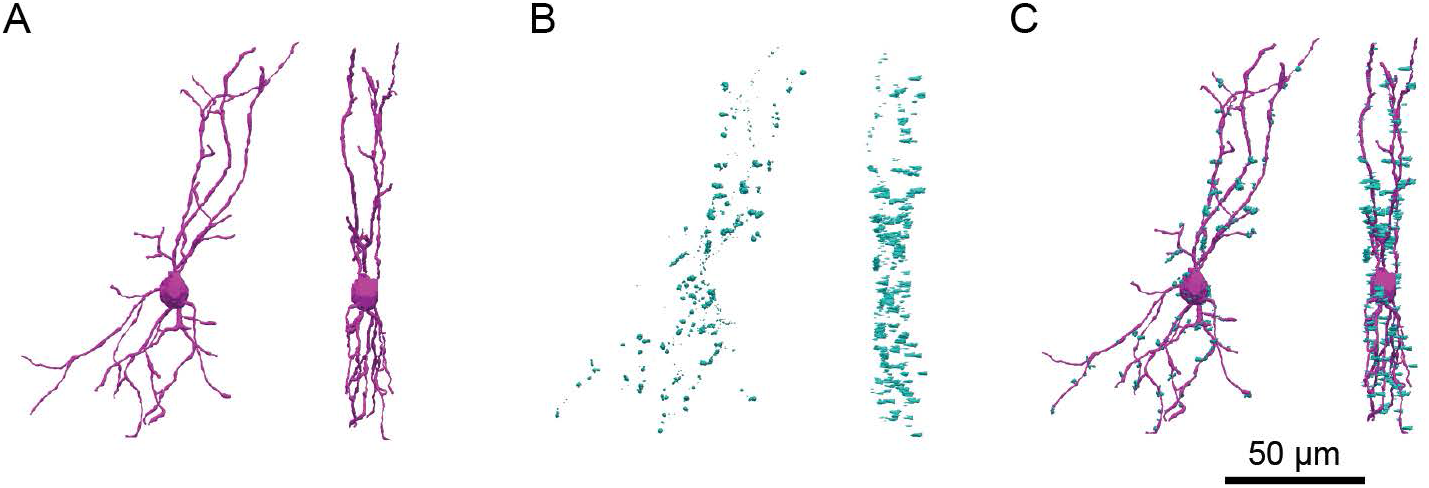
Example reconstruction of a biocytin-filled VIP neuron with closely apposed VAChT+ puncta. **A)** The soma and full dendritic arbor of a VIP neuron were reconstructed in three dimensions. **B)** VAChT+ puncta located within 1 μm of the reconstructed VIP neuron were detected and reconstructed. **C)** An overlay of A and B shows that putative cholinergic inputs were distributed along the full extent of the dendrites of the reconstructed VIP neuron. Scale bar in **C** also applies to all panels.

**Figure 7.**
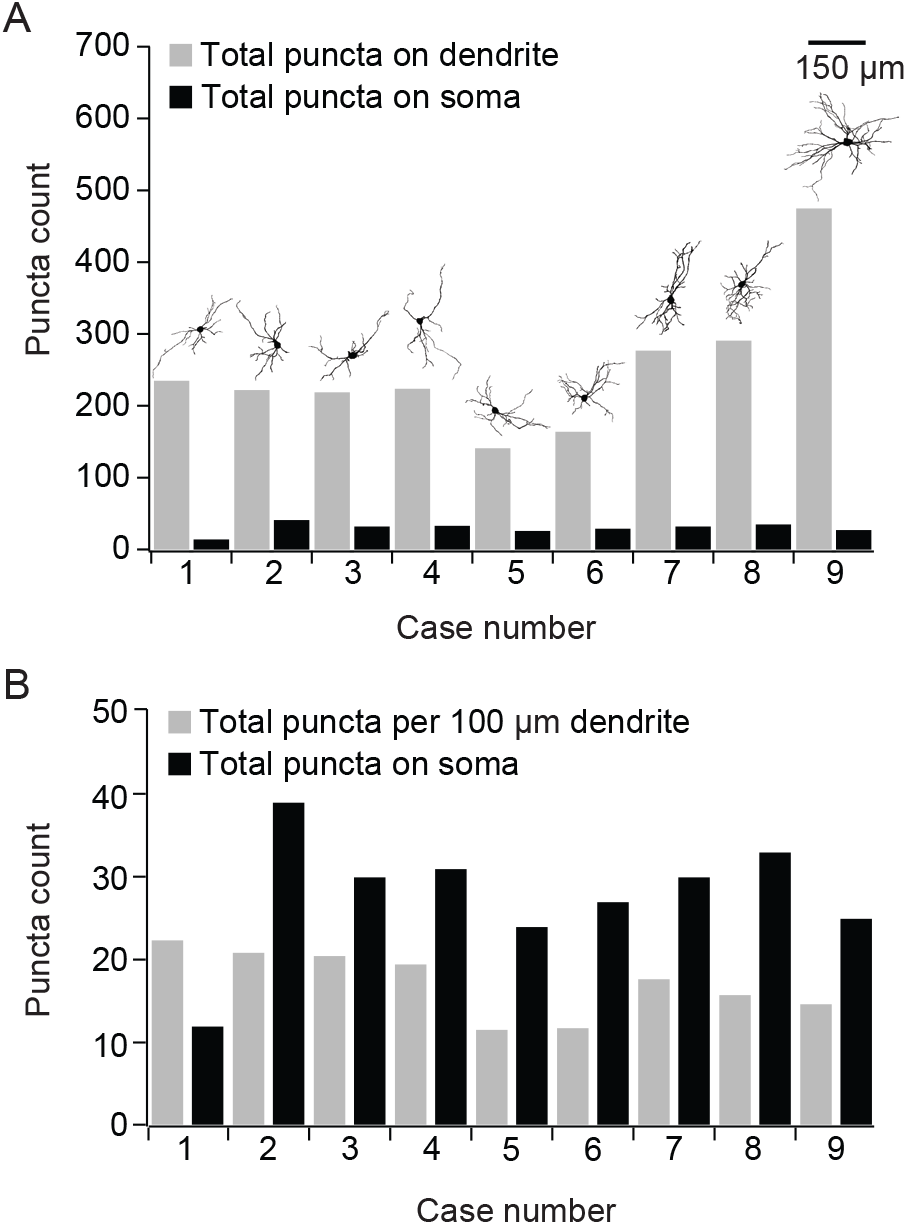
Fully reconstructed VIP neurons were lined with VAChT+ puncta throughout their dendritic arbors and along their somata. VIP neurons were filled with biocytin during whole-cell recordings, allowing subsequent reconstruction of their somata and their full dendritic arbors. **A)** Total VAChT+ puncta counts for the somata and dendritic arbors from 9 fully reconstructed VIP neurons. Insets above each pair of bars show the reconstruction of the VIP neuron that the data were taken from. The scale bar inset applies to the VIP neuron reconstructions. **B)** Same data as **A** but the dendritic puncta counts were normalized to 100 μm of dendritic length to facilitate comparisons neurons, which varied in their total dendritic length. Cases in **A,B** are arranged in order of increasing total dendritic length, with the neuron having the least total dendritic length on the left and the neuron with the most on the right.

The above results support the hypothesis that cholinergic inputs are common along the dendrites of VIP neurons in the ICc, but they do not address whether these inputs preferentially impinge upon a subregion of the dendritic arbor, for example, the distal dendrites, or are evenly distributed throughout the dendritic arbor. To distinguish between these possibilities, we performed a Sholl analysis on the reconstructed neuron data set. The Sholl analysis was conducted using nested virtual shells centered around the centroid of the soma of each reconstructed VIP neuron. Each shell was 10 μm thick, spherical in shape, and had a radius 10 μm greater than the previous shell (**Figure 8A,B**). For each shell, we determined the total dendritic length located within the thickness of the shell and the number of VAChT+ puncta closely apposed to dendrites located within the thickness of the shell. We then normalized the puncta count in each shell per 100 μm of dendritic length. The first Sholl shell (radius = 10 μm) was excluded from subsequent analysis because it primarily contained soma with little or no dendrite. The resulting data show that putative cholinergic inputs were rather evenly distributed along the length of VIP neuron dendrites (**Figure 8C**). The data reveal a tendency for a higher puncta count in the first Sholl shell, which is likely due to the inclusion of some somatic contacts in this region, but otherwise, the average puncta count per Sholl radius across cells (**Figure 8C**, black line) did not vary much from the grand average of 17 ± 4 puncta per 100 μm of dendrite that was calculated above (mean ± SD; **Figure 8C**, dashed line). Overall, these results suggest that putative cholinergic inputs are common along the entire length of the dendritic arbors of VIP neurons.

**Figure 8.**
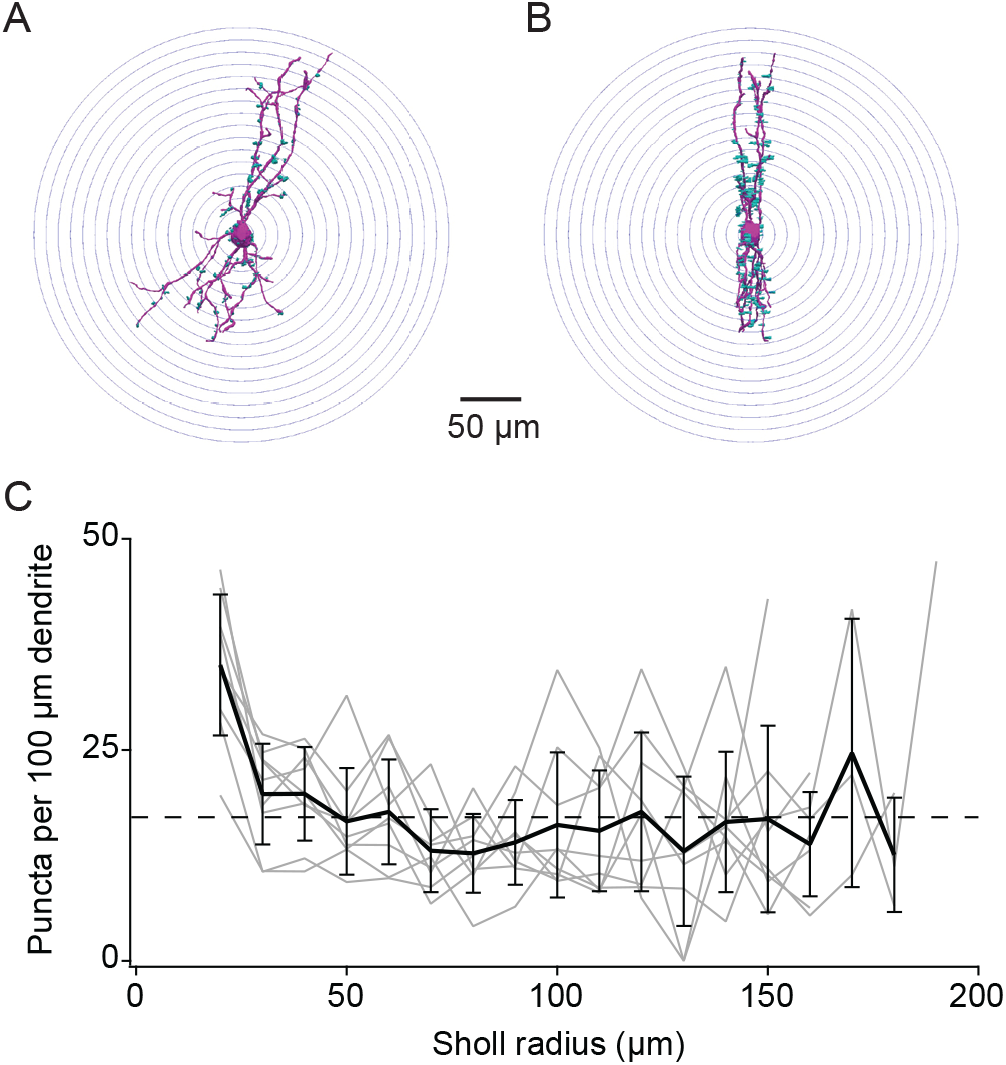
Sholl analysis shows that putative cholinergic inputs were relatively evenly distributed along the full length of VIP neuron dendritic arbors. **A,B)** Illustration of the Sholl analysis approach. Threedimensional, spherical shells of increasing diameter extended from the centroid of the soma of a reconstructed VIP neuron. Front and side views of the reconstructed neuron (magenta) and its associated VAChT+ puncta (cyan) are shown in **A** and **B**, respectively. **C)** Plot of the number of VAChT+ puncta detected in each Sholl shell starting with a Sholl radius of 20 μm. The first Sholl radius was excluded because it primarily encompassed somata and therefore contained little dendritic length. Gray lines represent data from individual cells. The solid black line shows the mean puncta count for each Sholl radius that contained data from 2 or more neurons. The dashed black line represents the overall average puncta count per 100 μm dendrite, which was 17. Error bars show standard deviation.

### VAChT+ puncta adjacent to dendritic spines

An important feature of IC VIP neurons is that their dendrites are studded with dendritic spines (Goyer et al. 2019). These spines likely represent sites of glutamatergic synaptic input, although that has yet to be confirmed. Interestingly, while analyzing the data from the fully reconstructed VIP neurons, we noticed that cholinergic puncta were occasionally located adjacent to the dendritic spines of VIP neurons in the ICc. **Figure 9** provides examples of three such cases where a VAChT+ punctum was located adjacent to a dendritic spine on a VIP neuron. Instances of cholinergic synapses onto dendritic spines have been reported in some ultrastructural studies (Smiley et al. 1997; Turrini et al. 2001), raising the possibility that cholinergic input to VIP neurons might sometimes occur at dendritic spines. However, a more complicated arrangement is also possible, in which the cholinergic bouton does not synapse onto the spine but rather uses cholinergic signaling to modulate the release probability of the glutamatergic bouton that might instead be synapsing onto the spine. It will be interesting to differentiate between these possibilities in future ultrastructural and physiological studies.

**Figure 9.**
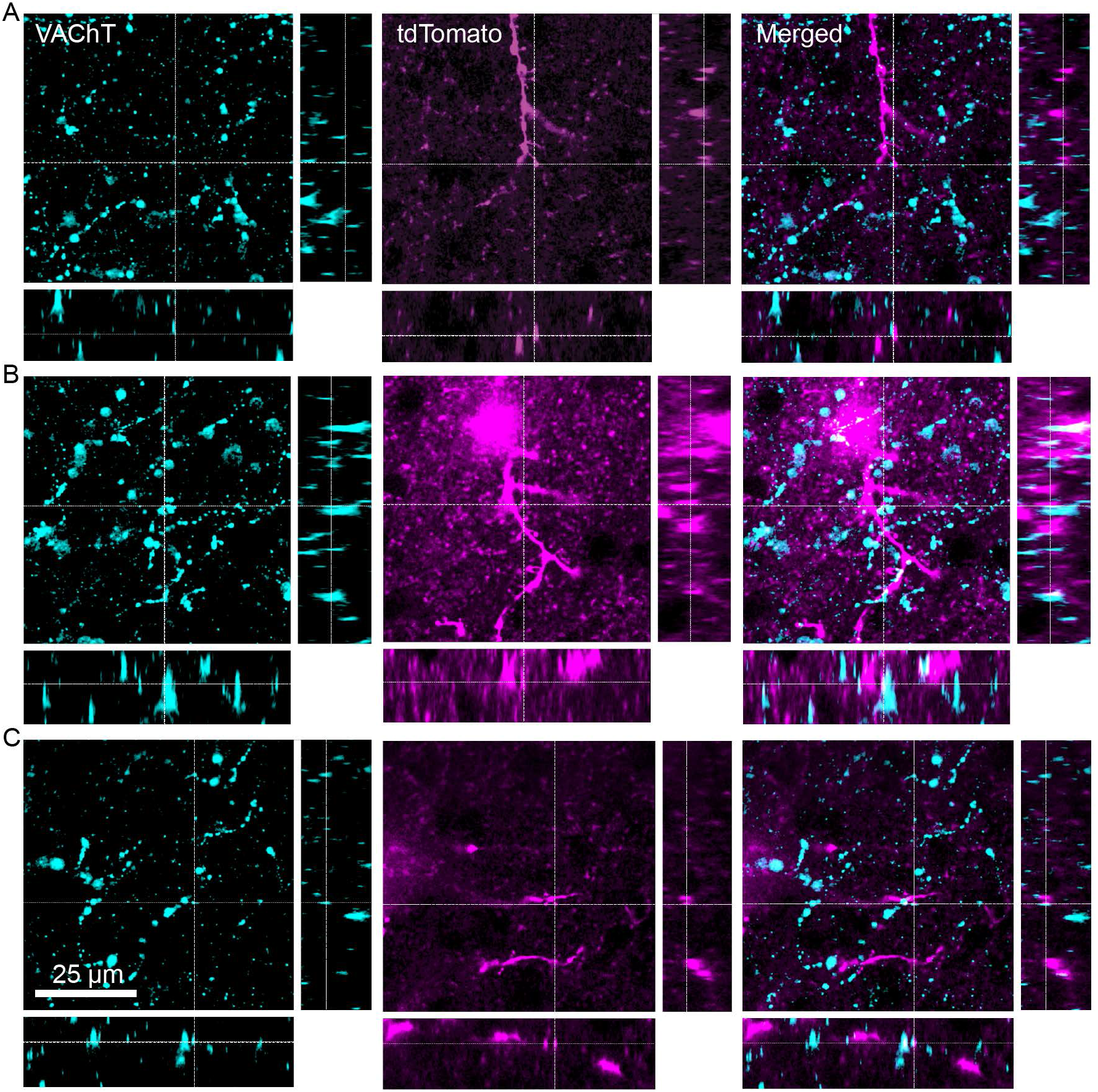
Cholinergic puncta were often observed in close apposition to the dendritic spines of VIP neurons. **A-C** show examples of VAChT+ puncta located directly adjacent to dendritic spines from three different fully reconstructed VIP neurons. The color scheme and organization of the figure is the same as described in the Figure 3 legend. The scale bar in **C** applies to all panels. Data are from 3 mice.

## Discussion

The data presented here provide multiple lines of anatomical evidence supporting the hypothesis that VIP neurons in the IC are subject to modulation by cholinergic signaling. Using fluorescence in situ hybridization, we found that 98% of VIP neurons express mRNA encoding α_3_ nAChR subunits, and 61% express mRNA for both α_3_ and β_4_ nAChR subunits. We also found that α_3_ and β_4_ nAChR subunits were expressed in other cells throughout the IC, pointing to a broad role for α_3_- and β_4_-containing nAChRs in the IC. We examined the distribution of putative cholinergic release sites on VIP neurons in the ICc using two approaches: one assaying the distribution of VAChT+ puncta adjacent to isolated VIP neuron somata and dendritic segments, the other assaying the distribution of VAChT+ puncta adjacent to the somatic and dendritic arbors of fully reconstructed VIP neurons. Both approaches revealed that presumptive cholinergic boutons and terminals are commonly found adjacent to the somata and dendrites of VIP neurons, while the latter approach also showed that cholinergic puncta were relatively evenly distributed along the length of the dendritic arbors of VIP neurons. In addition, we observed that cholinergic puncta could be located adjacent to VIP neuron dendritic spines, where they might directly synapse onto VIP neurons or influence presynaptic release from glutamatergic synapses that instead contact those spines. Overall, these results suggest that cholinergic signaling is positioned to exert widespread influence over computations in VIP neurons, likely shaping synaptic integration throughout the dendrites and the soma.

### nAChR subunits in the IC

Previous in situ hybridization and binding studies indicated that α_3_, α_4_, α_7_, β_2_, and β_4_ nAChR subunits are expressed in the IC, but it has remained unclear whether specific populations of IC neurons express subsets of these (Clarke et al. 1985; Wada et al. 1989; Morley and Happe 2000; Whiteaker et al. 2002; Marks et al. 2002; Salas et al. 2003; Gahring et al. 2004; Happe and Morley 2004; Marks et al. 2006; Bieszczad et al. 2012; Sottile et al. 2017). The in situ hybridization results reported here show that nearly all VIP neurons expressed mRNA for α_3_ nAChR subunits, and 61% expressed mRNAs for both α_3_ and β_4_ nAChR subunits. Interestingly, this indicates that while most VIP neurons express α_3_β_4_* nAChRs, 37% of VIP neurons express α_3_ nAChR mRNA but not β_4_ nAChR mRNA. Thus, in a minority of VIP neurons, the α_3_ subunit presumably forms functional nAChRs by pairing with non-β_4_ subunits. This extends the interpretation of our recent pharmacological study, which showed that cholinergic responses in VIP neurons were largely blocked by SR16584 (Rivera-Perez et al. 2021), an antagonist reported to be selective for α_3_β_4_* nAChRs (Zaveri et al. 2010). A likely explanation is that SR16584 blocks not only α_3_β_4_* nAChRs but also other α_3_-containing nAChRs. Indeed, the study that identified SR16584 as an α_3_β_4_*-selective antagonist compared drug effects across only three types of nAChRs: α_3_β_4_, α4β2, and α7 (Zaveri et al. 2010).

Based on results from our previous pharmacological study and the present study, we therefore propose that nearly all VIP neurons express α_3_-containing nAChRs and that the majority of VIP neurons express α_3_β_4_* nAChRs (Rivera-Perez et al. 2021). This parallels results from another recent in situ hybridization study, which reported that putative glutamatergic neurons in the IC could express α4 and β2 subunits together, β2 alone, or neither subunit (Sottile et al. 2017). In future studies, it will be important to further define the combinations of nAChR subunits expressed in distinct populations of IC neurons and to determine whether the expression of specific combinations of nAChR subunits points to different functional roles for cholinergic signaling in different populations of IC neurons. Since α_3_- expressing VIP neurons can be divided into subpopulations that either express or do not express β_4_ subunits, VIP neurons are a prime candidate for such functional studies.

### Technical considerations

Our results show that the somata and dendrites of VIP neurons in the ICc were closely adjacent to numerous VAChT+ puncta. Since these observations were made using light microscopy, we cannot be certain that these VAChT+ puncta represented cholinergic release sites, but several lines of evidence suggest that they were. First, it is well established from electron microscopy studies that VAChT immunoreactivity occurs in subcellular domains of cholinergic neurons where concentrated pools of vesicles are present (Gilmor et al. 1996; Weihe et al. 1996; Turrini et al. 2001). The IC is not known to contain cholinergic neurons, and thus the presence of punctate VAChT labeling in the IC is consistent with local concentrations of acetylcholine-containing vesicles in cholinergic axons, most of which presumably originated in the PPT and LDT (Motts and Schofield 2009; Noftz et al. 2020).

Second, acetylcholine release is thought to occur in two modes: through conventional synaptic transmission, where a presynaptic bouton is separated from a postsynaptic density by a nanometer-scale gap, and through volume transmission, where neurotransmitter released into the extracellular space diffuses a micron-scale distance to reach postsynaptic receptors. There is significant controversy about the preponderance of each of these modes of cholinergic transmission, but it is now generally accepted that cholinergic signaling in many brain regions is mediated by a combination of synaptic and volume transmission (see recent Dual Perspectives by: Disney and Higley 2020; Sarter and Lustig 2020). Whether cholinergic transmission in the IC occurs through synaptic or volume transmission or both remains unclear, and there is a pressing need for ultrastructural studies in the IC to examine this question in detail. With respect to the present study, our observation that an average of 12-17 VAChT+ puncta were located within 2 μm of each 100 μm of VIP neuron dendrite suggests, at minimum, ample opportunity for release of acetylcholine onto VIP neurons through volume transmission. However, if the IC is similar to other brain regions, it is quite likely that a portion of the VAChT+ puncta we observed represent conventional synapses.

Third, we previously observed that exogenous applications of acetylcholine elicited depolarization and firing in 92% of tested VIP neurons (n = 116 out of 126) and that this response was mediated by nAChRs (Rivera-Perez et al. 2021). It seems unlikely that VIP neurons would express nAChRs and exhibit strong responses to acetylcholine if there were not a significant endogenous source of cholinergic transmission to VIP neurons. Furthermore, since the α_3_β_4_* nAChRs expressed by most VIP neurons have a relatively high affinity for acetylcholine (Krashia et al. 2010), as well as long single-channel open times and burst durations (David et al. 2010), they are good candidates for activation by acetylcholine released either synaptically or through volume transmission. Considering the above arguments, we conclude that the immunofluorescence data reported here point to a strong probability that VIP neurons in the ICc receive a meaningful amount of cholinergic input to their somata and throughout their dendritic arbors. Since our in situ hybridization results indicated the VIP neurons in the dorsal cortex and lateral cortex of the IC also expressed α_3_ and often β_4_ nAChR subunit mRNA, it is likely that VIP neurons in these IC shell regions also receive meaningful amounts of cholinergic input, although that remains to be directly tested.

### Functional implications for cholinergic signaling to VIP neurons

Our finding that putative sites of cholinergic input are present throughout the dendritic arbors of VIP neurons in the ICc compliments the observation that cholinergic axons from the PPT project throughout the IC without apparent bias for specific IC subdivisions or domains within subdivisions (Noftz et al. 2020). Both results point to a broad role for cholinergic signaling in changing the state of processing in the IC, as opposed to a more targeted role that would selectively influence specific subsets of computations. Indeed, while it is not yet known whether specific sources of non-cholinergic input target specific subdomains of VIP neurons or are distributed more homogenously, our results suggest that cholinergic input is situated to broadly influence synaptic integration in VIP neurons in either scenario. This contrasts, for example, with the situation in Layer 5 pyramidal neurons in neocortex, where cholinergic input to the apical dendrites dramatically enhances excitability while cholinergic input to the soma and axon has little effect (Williams and Fletcher 2018). It will be interesting to see whether other classes of IC neurons receive similarly large and distributed patterns of cholinergic input or if cholinergic signaling can have more targeted effects in certain neurons. As a class of stellate neurons, it is possible that VIP neurons interact with cholinergic input differently than the disc-shaped cells that make up the majority of neurons in the ICc.

In the present study, we were unable to determine whether putative cholinergic boutons originated from the PPT, LDT, or another cholinergic structure. It therefore remains possible that the sources of cholinergic boutons contacting VIP neurons vary as a function of location within the somatodendritic domain of individual VIP neurons or location of VIP neurons within the ICc. Importantly, we focused here only on VIP neurons located in the ICc, and the situation might be different in the dorsal and lateral cortices of the IC. In addition, cholinergic neurons in the rostral PPT have different functional roles than those in the caudal PPT (Mena-Segovia and Bolam 2017), providing another potential avenue for more nuanced modulation of VIP neurons by cholinergic input. While it will be important to sort out these details, the present results combined with our recent pharmacological results (Rivera-Perez et al. 2021) suggest that VIP neurons possess the anatomical and physiological features expected of cells that are subject to extensive modulation by cholinergic input. Since VIP neurons are glutamatergic neurons that project broadly within the IC and to several long-range targets, including the auditory thalamus and periaqueductal gray, we therefore propose that behavioral state changes signaled by cholinergic input to the IC might be broadcast to an array of brain regions by broadly enhancing the excitability of VIP neurons.

## Acknowledgements

We thank Nichole Beebe, Jeffrey Mellott, William Noftz, and Brett Schofield for helpful advice and feedback, and Marina Silveira for help with the in situ hybridization experiments. This work was supported by National Institutes of Health Grants F31 DC019292 (LMR-P) and R01 DC018284 (MTR).

## Statements and Declarations

### Competing Interests

The authors declare no competing financial or non-financial interests.

## Author Contributions

All authors conceived of the study, designed experiments, prepared figures, composed and revised the manuscript, and approved the submitted version. JTK and LMR-P performed experiments and analyzed data. MTR obtained funding and supervised the study.

## Notes

### Competing Interest Statement

The authors have declared no competing interest.

